# Wirelessly-Powered Ingestible Electronic Capsule for Non-invasive Gastrointestinal Optogenetics

**DOI:** 10.1101/2024.08.30.610532

**Authors:** Mohamed Elsherif, Rawan Badr El-Din, Zhansaya Makhambetova, Heba Naser, Maylis Boitet, Rahul Singh, Keonghwan Oh, Revathi Sukesan, Sohmyung Ha, Khalil B. Ramadi

## Abstract

Optogenetics enables the activation and inhibition of neurons with cell specificity. The gut harbors intricate networks of enteric and central neurons. Uncovering these neuronal pathways in vivo is challenging with traditional neuroscience probes due to the highly motile and harsh gut environment. Here we report the development of an ingestible electronic capsule for non-invasive optical gut stimulation (ICOPS) in rodents. ICOPS is powered wirelessly via a transmitter coil, dosed via oral gavage, and safely excreted without causing obstruction. ICOPS permits modular interchangeability of onboard light-emitting diodes (LEDs) for illumination. We exemplify this with optical irradiance at 470 nm, a commonly-used wavelength in optogenetics for activating channelrhodopsin2. ICOPS features a micro-LED (µLED), a 460-turn coil wound around a ferrite core, and a resonating capacitor. We optimized the transmitting and receiving circuits to achieve maximum power transfer at low operating frequencies (45-140 kHz), overcoming challenges like loose coupling and misalignment. The capsule operates effectively at a distance up to 12 cm longitudinally, 9 cm laterally, and 75° rotational angle relative to the magnetic field. Specific absorption rate (SAR) calculations indicate transmitter-induced SAR levels within safe limits for the occupational environment at 6 A_rms_ and 45 and 63 kHz frequencies ICOPS is robust and transits through the rat gastrointestinal (GI) tract in under 20 hours intact. We demonstrate in vivo functionality and viability of ICOPS using IVIS micro-computed tomography (µCT). ICOPS could pave the way for non-invasive optogenetic interfacing of enteric neural circuits towards their use to regulate motility, visceral pain, and other gastrointestinal disorders.

## Introduction

The gastrointestinal tract (GIT) harbors the enteric nervous system (ENS), the body’s second-largest network of neurons after the brain.^1^ The ENS consists of intrinsic sensory neurons, interneurons, and excitatory and inhibitory motor neurons within the gut wall. This neural network forms ganglionated networks encircling the gut.^2^ Significant efforts have been devoted to classifying enteric neurons based on their axonal projections, morphology, and electrophysiological characteristics, providing insights into their functional roles as sensory, inter, and motor neurons. Within specific regions of the gut and across species, classes of enteric neurons exhibit consistent combinations of proteins, neurotransmitters, and other neurochemicals, forming what is termed their ‘neurochemical code’.^3^ While traditionally used for classification and identification purposes, the neurochemical coding of enteric neurons now holds promise for neurogenetic targeting, allowing for the selective expression of light-sensitive ion channels in specific classes of enteric neurons.^4^ Consequently, the presence of the ENS in the gut wall presents a unique opportunity to precisely control the activity of targeted neuron classes using optogenetics, leveraging their neurochemical coding for precise temporal activation or inhibition.

Current therapeutic approaches for enteric neuromodulation, such as nonspecific electrical stimulation, indiscriminately activate both sensory and motor axons.^5^ Similarly, chemical stimulation is challenging to target spatially, and systemic transport inevitably leads to a broad spectrum of effects across multiple organ systems.^6^ Technologies enabling non-invasive and remote stimulation of specific neurons in the GI tract have long been sought after for experimental investigations into neural systems and for the clinical treatment of GI and psychiatric disorders.^7^ With its high spatial, temporal, and cell-type specificity, optogenetic stimulation could be used for understanding the enteric neural pathways that regulate pain and motility, towards the development of therapies to address various GI disorders.

Optogenetics enables activation or inhibition of specific cells by transfecting with light-gated ion channels, followed by illuminating with light of specific wavelengths to which ion channels are sensitive.^8^ The most commonly used light-gated channel is channelrhodopsin-2, which has a response peak at 470 nm light.^9,10^ Physiologic tissue absorbs and scatters blue light, rendering it challenging to deliver externally.^11^ Despite the development of red-shifted variants of rhodopsins (light-gated ion channels), their action spectra still do not fully align with the near-infrared (NIR) optical window (ranging from 650 to 1350 nm), where light achieves its maximum depth of penetration in biological tissues.^12–16^

As a result, optical illumination for channelrhodopsin activation requires the insertion of an optical fiber or micro-sized light-emitting diode (μLED). Such an approach is well established in the brain where probes can be implanted and secured to the skull, but more challenging in soft tissues such as the GI tract.^17^ Existing wireless technologies designed for optogenetic control of the brain and spinal cord are efficient but prove unsuitable for stimulating the GI tract.^17^ For instance, devices incorporating serpentine interconnect structures and microfabricated features function effectively in brain applications but fail reliably in the gastrointestinal environment due to mechanical issues with metal features and encapsulating layers, exacerbated by the natural movements and muscle contractions within the gut.^18^

Here we introduce a non-invasive platform for targeted optical illumination within the GI tract of rats. Our ingestible electronic capsule for non-invasive optical gut stimulation (ICOPS) is equipped with micro-sized light-emitting diodes (µLEDs) and electronics for wireless powering based on magnetic induction. ICOPS bypasses traditional semiconductor fabrication methods, opting instead for micro-resolution 3D printing to produce ICOPS. It provides ample light intensity across large spatial fields to reliably activate the GI tract without obstruction. This technology offers unique opportunities for noninvasive, wireless control of the ENS and gut-brain axis enabling stimulation at multiple times and locations on-demand.

## Results

We first set out to develop a device that could be ingested by rats and deliver optical illumination within the gut (Figure 1 A-C). We designed ICOPS with dimensions of 2.7 mm in diameter and 18 mm in length, weighing 0.22 g to ensure compatibility with the largest ingestible capsule size for rats (9el capsule, 2.7 mm in diameter, and 23 mm in length) (Figure 1D). We selected a μLED emitting predominantly at 460 nm, measuring 240×320 µm, small enough to fit within a 9el capsule, and capable of activating channelrhodopsin-2 (Figure 2 A and B). ICOPS incorporates materials for wireless powering via magnetic induction, a 460-turn coil wound on a ferrite core, a μLED, and a resonant capacitor (Figure 1 C).

**Figure 1.**
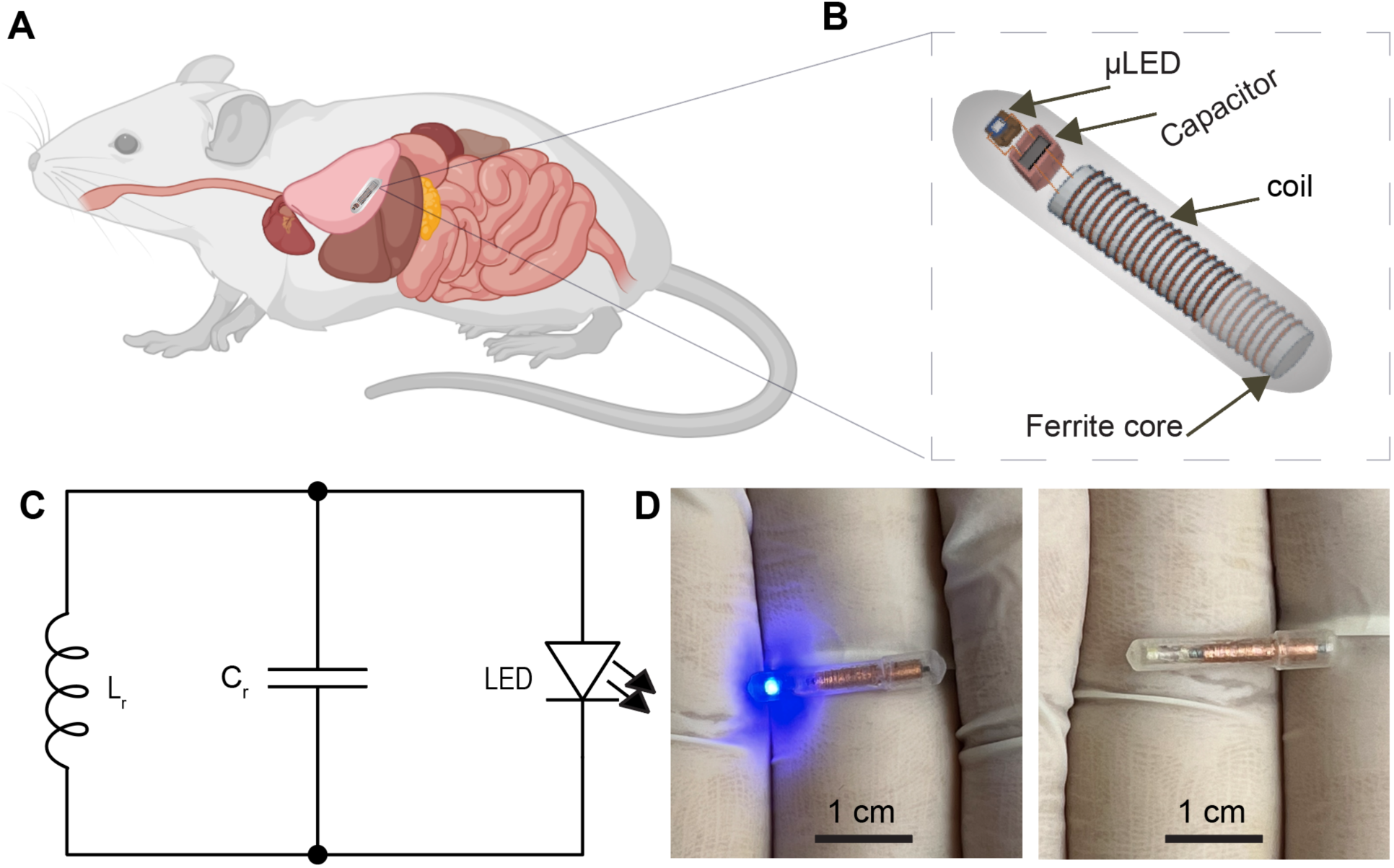
Overview of the ingestible electronic capsule (ICOPS). (A-B) Schematic representation of the capsule administered to a rat phantom. (C) Diagram illustrating ICOPS circuit. (D) Photographs depicting ICOPS in its On and Off conditions.

**Figure 2.**
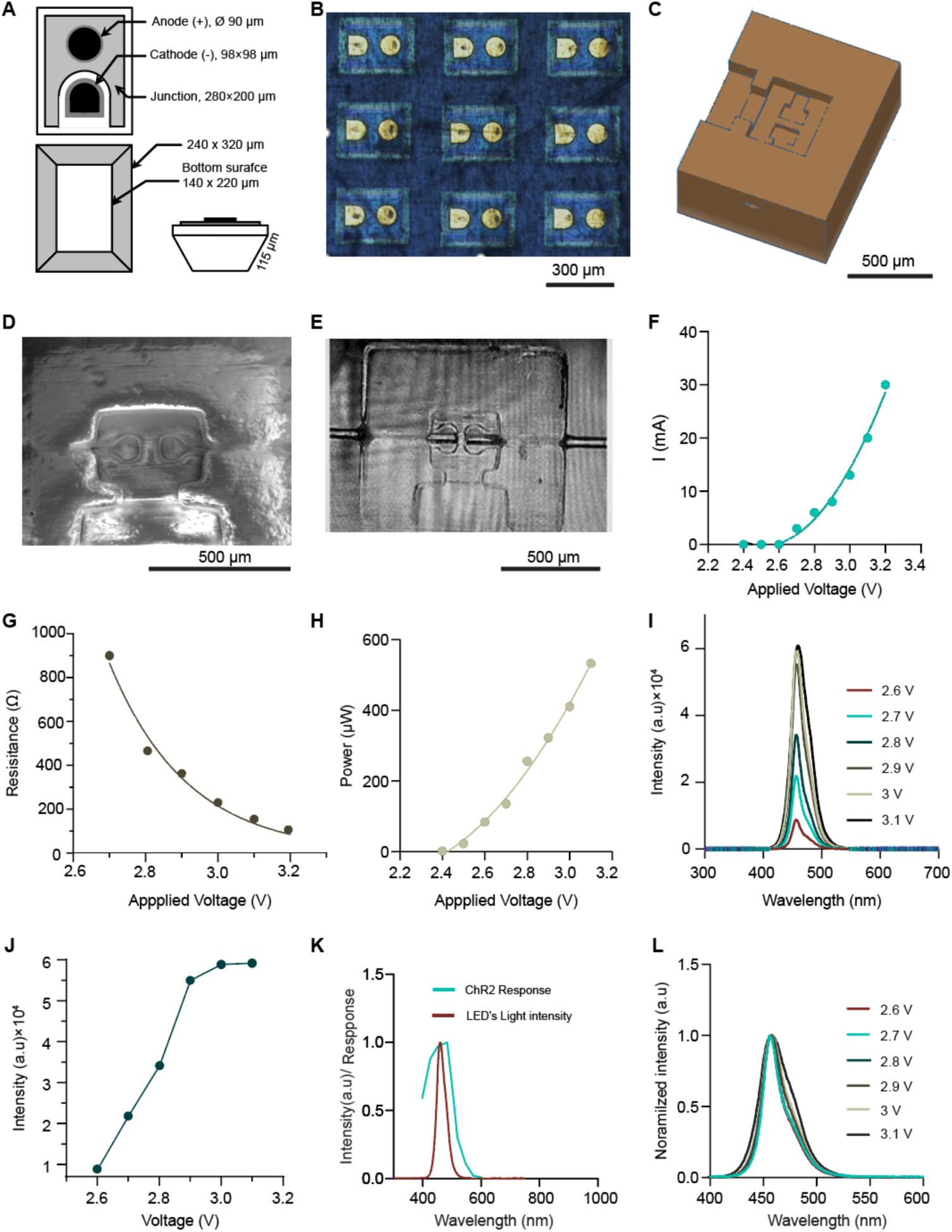
(A) Diagram illustrating the blue μLED, showing its top, bottom, and side views. (B) (Optical microscope image of a batch of the blue μLEDs. (C) Schematic of the mold designed to connect the μLED. (D) SEM image of the 3D printed μLED mold. (E) Optical microscope image of the mold with wires threaded to connect the μLED. (F) I-V characteristic curve of the blue μLED. (G) Resistance of the μLED versus applied voltage. (H) Optical power emitted from the μLED under varying applied voltages. (I) Light intensity of the μLED versus wavelength under different applied voltages. (J) Light intensity of the blue light emitted from the μLED versus applied voltage. (K) Comparison of the μLED light intensity to the ChR2 response. (L) Normalized light intensity versus wavelength under different applied voltages.

### Optical and Electrical Characteristics of the Selected μLED

We developed our system to facilitate component interchangeability. Rather than relying on microfabrication techniques tailored for the specific μLED, we employed 3D printing to create a mold for μLED integration. This mold includes channels for micro-wire threading, traps for soldering epoxy, and housing for μLED attachment (Figure 2 C-E). The 3D printing method enables efficient μLED replacement without the time and complexity associated with optimizing semiconductor fabrication stages and designs.

To assess the electrical characteristics of the μLED, we investigated its current-voltage characteristic curve (I-V curve). We mapped µLED performance over an operational voltage range of 2.4 V to 3.2 V (Figure 2F). The µLED begins to emit light at 2.7 V, with the current reaching the maximum forward current (30 mA) at 3.2 V. The I-V curve demonstrates that the operation voltage range of the µLED is in the range of 2.7-3.2 V. Based on the I-V curve, we determined the μLED resistance to be within the range of 90-900 Ω (Figure 2G).

To effectively stimulate ChR2 with blue light, the irradiance must reach at least 1 nW/mm^2^.^19,20^ Accordingly, we measured the optical power of the μLED across different applied voltages within its operational range of 2.7-3.2 V (Figure 2H) and observed a nonlinear relationship mimicking I-V curve, with power reaching 600 µW or 8.7 µW/mm^2^ at an applied voltage of 3.1 V. Notably, even at the minimum operation voltage of 2.7 V, the μLED output power exceeded the required irradiance, reaching 300 μW or 3 µW/mm^2^. Additionally, we measured the light intensity of the µLED within its operational voltage range to examine the dominant wavelength and bandwidth of the emitted blue light (Figure 2 I and J). The collected spectra indicate that the dominant wavelength is 460 nm, aligning with the response of ChR2 (Figure 2K). Additionally, the full width at half maximum (FWHM) was observed to increase within the operational range, ranging from 25 to 30 nm with increasing applied voltages (Figure 2L).

### Wireless Powering the Capsule

The majority of ingestible electronic devices rely on onboard coin cell or silver oxide batteries. However, these are too large to be safely ingested by rodents within 9el capsules. To address this challenge, we developed a wireless powering system based on magnetic induction. This approach utilizes a primary-powered coil as a transmitter and a secondary coil as a receiver.^21^ A waveform generator supplies an electric signal to an RF amplifier, which amplifies the signal before transmitting it through the primary coil, connected in series with a resonating capacitor (Figure 3 A and B). The external transmitter coil, wound from 18-gauge Litz wire, comprises 50 turns with a diameter of 6.8 cm and a length of 7 cm, suitable for rats weighing 250-750 g. The transmitter operates at a low-frequency range of 45-140 kHz to minimize tissue absorption and reduce the risk of heating from high-frequency magnetic fields, which are more readily absorbed by physiological tissues.^22^

**Figure 3.**
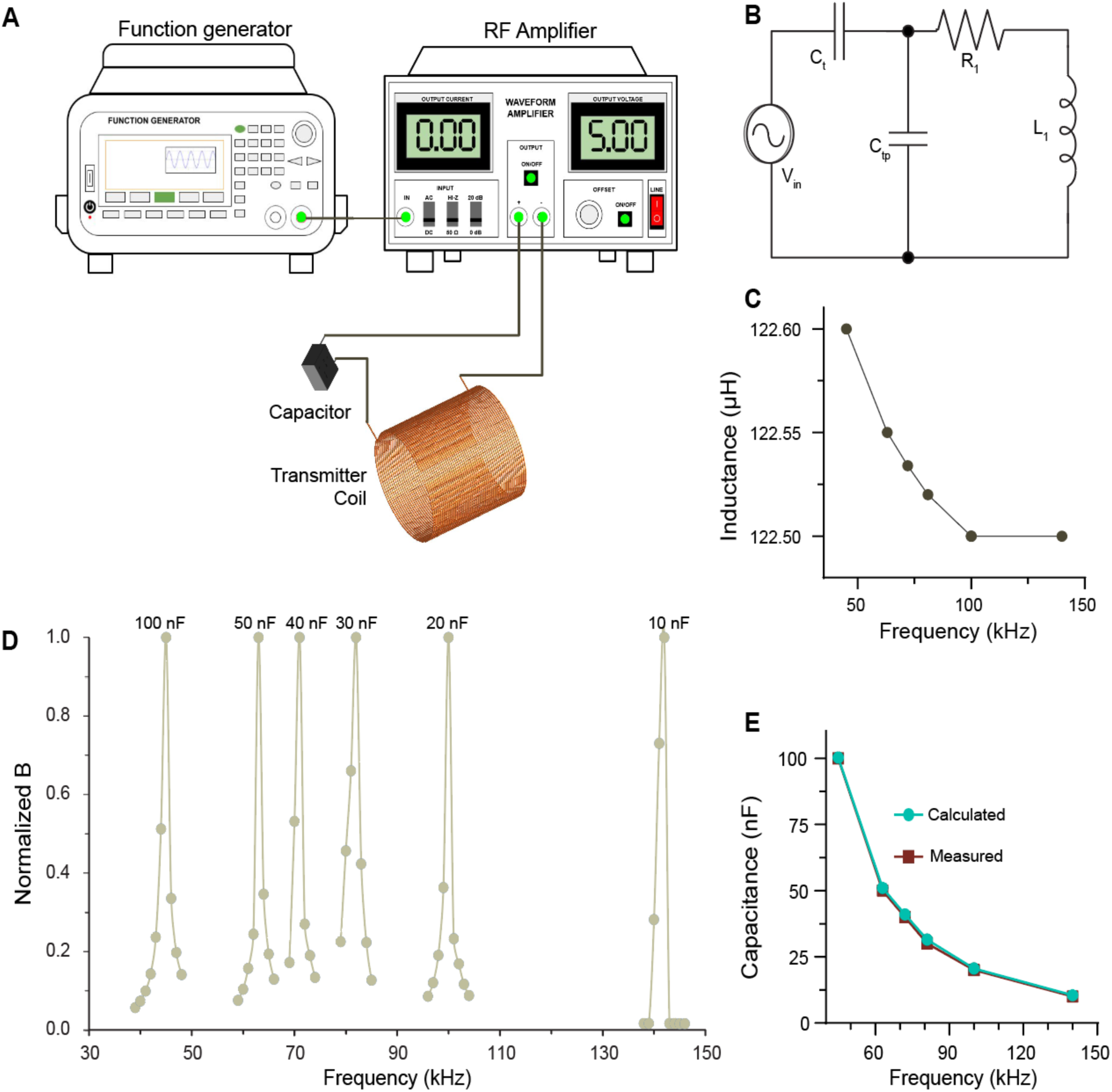
The transmitter circuit used for powering ICOPS. (A) Schematic diagram of the transmitter circuit. (B) Diagram of the transmitter circuit. (C) Measured inductance of the transmitter coil. (D) Normalized measured magnetic flux density versus sweep frequency with capacitors of 10, 20, 30, 40, 50, and 100 nF resonating the circuit. (E) Calculated capacitance of the transmitter coil across the operating frequency range, compared to the resonant capacitance of the transmitter extracted based on magnetic flux density measurements.

Inductor impedance is known to linearly increase with frequency up to the resonance frequency.^23^ To overcome this, we used capacitive resonance to minimize the impedance of the transmitter coil.^24^ This allows us to drive a high current through the transmitter coil using a low-output power amplifier, thereby generating a sufficiently strong magnetic field to power the remote capsule. Capacitive reactance effectively reduces the inductor’s impedance by matching with the inductor reactance at a specific operating frequency, ensuring efficient power transfer. However, the resonance capacitive technique requires a capacitor of a specified capacitance to be connected to the transmitter circuit for each operating frequency.

To resonate the transmitter circuit, we initially measured the inductance of the transmitter coil using an LCR meter across the operating frequency range of 45-140 kHz (Figure C). The measured inductance ranged from 122.5 to 122.6 µH, with only a minimal variation of 0.1 µH within this frequency range. These values were crucial for calculating the resonance capacitance of the transmitter circuit using the formula:

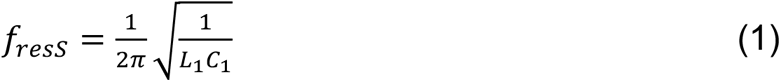

where *f*_resS_ is the resonance frequency, L_1_, and C_1_ are the self-inductance and the resonant capacitance of the transmitter, respectively.^25^ Figure 3E shows the calculated resonating capacitances for several operating frequencies.

To verify optimal performance, we individually connected capacitors with capacitance similar to calculated values in series for resonating the transmitter circuit. We monitored the magnetic flux density (*B*) generated in the transmitter coil while sweeping the operating frequency around the expected circuit resonance frequency for each capacitor (Figure 3D). The magnetic flux density remained relatively low across the frequency spectrum and peaked at the expected resonance frequencies of 45, 63, 71, 82, 100, and 140 kHz, corresponding to the resonating capacitances of 100, 50, 40, 30, 20, and 10 nF, respectively, confirming their effectiveness within the desired operating frequency range (Figure 3E). The relationship between the operating frequency of the transmitter circuit and the resonant capacitance is inversely proportional and nonlinear, as illustrated in Figure 3E. Furthermore, we measured the parasitic capacitance of the transmitter coil in the operating frequency range, which matched the required resonating capacitance for the transmitter circuit to neutralize its inductance (Figure S1A).

We used a receiver coil within the capsule to harvest the magnetic energy powering the µLED, with a capacitor employed to tune the circuit (Figure 4A). The receiver coil, measuring 2 mm in diameter and 10 mm in length, was fabricated to fit within the 9el capsule size. A ferrite core was used in the receiver coil to increase its induced peak voltage. Ferrite cores have also been shown to enhance the transmission efficiency in loose coupling, and lateral and angular misalignment conditions.^26^

**Figure 4.**
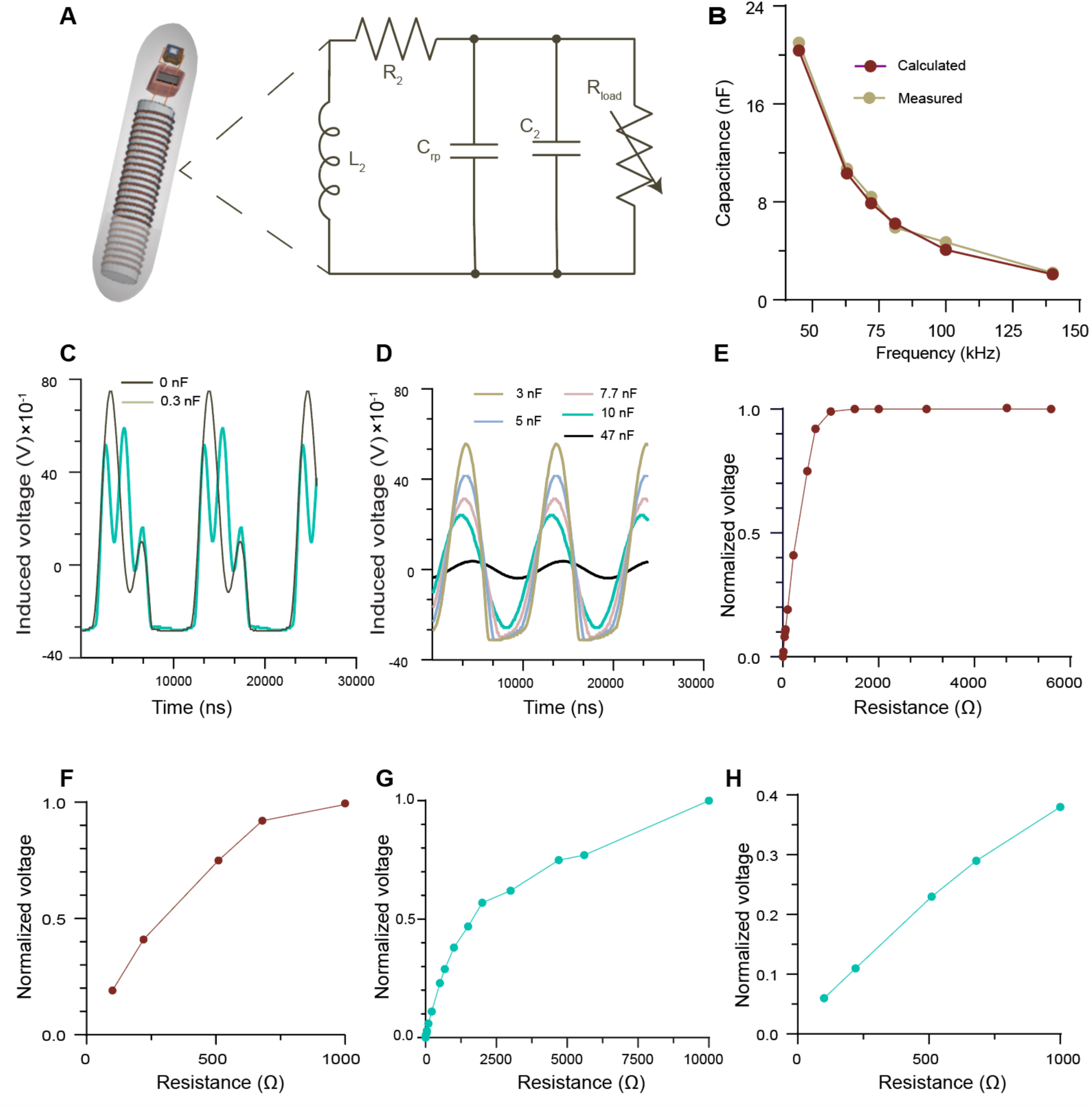
(A) Schematic of ICOPS and diagram of its electric circuit. (B) Resonating capacitance versus operating frequency of the receiver circuit extracted from measured μLED output power and compared with the calculated ones based on the self-inductance values. (C-D) Influence of the shunt capacitor on the induced voltage signal in the receiver circuit. (E-F) Induced voltage in the receiver circuit under varying load resistances at high input power in the transmitter circuit, operating frequency: 63 kHz. (G-H) Induced voltage in the receiver circuit versus load resistance at less input power in the transmitter circuit, operating frequency: 63 kHz.

We measured the self-inductance of the receiver coil using an LCR meter across the operating frequency range of 45-140 kHz, averaging approximately 626.5 μH (Figure S1B). The trend of inductance versus frequency differs from that of the transmitter coil, likely due to the ferrite core influence in this setup. Based on these inductance measurements, the resonant capacitance was calculated for each operating frequency using the formula:

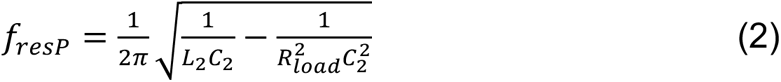

where *f*_resP_ represents the resonant frequency, *L_2_* is the self-inductance of the receiver, *C_2_* is the capacitance, and *R_loa_*_d_ is load resistance (assumed to be 900 Ω, the resistance of the µLED at 2.7 V applied voltage).^25^ We calculated resonant capacitances at the operating frequencies of 45, 63, 71, 82, 100, and 140 kHz to be 20.3, 10.3, 7.9, 6.2, 4.1, and 2.1 nF, respectively (Figure 4B). To verify these values as the resonant capacitances, we connected the µLED to the receiver coil with the capacitor, adjusting the input voltage in the transmitter circuit to a minimum level (10-15 mV_p_) sufficient to illuminate the µLED only when the resonant capacitor was connected. We observed no illumination occurred when capacitors of higher or lower capacitance were used and the measured capacitances closely matched the calculated resonant values (Figure 4B). The test results revealed resonant capacitances of 21, 10.7, 8.4, 5.9, 4.7, and 2.2 nF at operating frequencies 45, 63, 71, 82, 100, and 140 kHz, respectively. Similar to the transmitter circuit, the relationship between resonant capacitance and operating frequency in the receiver circuit was found to be inversely proportional and nonlinear. Additionally, the parasitic capacitance of the receiver coil, measured using the LCR meter across the frequency range of 45-140 kHz, aligned closely with the calculated resonant capacitance of the transmitter circuit (Figure S1C), yielding values of 20.4, 10.3, 7.9, 6.2, 4.1, and 2.1 nF at operating frequencies of 45, 63, 72, 81, 100, and 140 kHz, respectively.

To investigate the impact of the tuning capacitor on the electrical signal generated by the receiver coil connected to the μLED, we applied a sinusoidal wave to the transmitter coil and monitored the induced voltage amplitude using an oscilloscope. Integrating the μLED into the circuit caused partial rectification of the electrical signal because the μLED shares characteristics with a PN junction diode, allowing current flow in the forward direction while blocking it in the reverse direction. Without a capacitor in the receiver circuit, the induced voltage signal exhibited rectification and distortion, characterized by multiple ripples (Figure 4C). Introducing a capacitor with a capacitance lower than the required resonating value reduced the number of ripples (Figure 4C). Using a capacitor having resonant capacitance completely eliminated the ripples, while higher capacitance capacitors decreased the amplitude of the generated signal but maintained the rectified waveform (Figure 4D). However, very high capacitance negated the rectification effect of the μLED and further decreased the signal’s amplitude (Figure 4D). Based on these measurements, we conclude that the capacitor in the receiver circuit serves a dual purpose: tuning the circuit and facilitating signal rectification by compensating for the parasitic capacitance of the μLED.^27^

### Variant Load and Tuning Capacitor Effects

Given that the μLED connected to the receiver circuit exhibits varying loads within the range of 100-900 Ω, and considering equation (2), alterations in the load resistance impact the resonant capacitance. Therefore, we measured the capacitance of the receiver coil connected in parallel with different loads in the range of 100-900 Ω while the system operated at 63 kHz. It was observed that the load resistance slightly influenced the capacitance of the receiver circuit, causing it to vary from 10.6 to 11.2 nF, a shift of 0.06 nF (Figure S1D).

We next examined the effect of the variable loads of the μLED on the amplitude of the induced voltage while both transmitter and receiver were tuned to operate at 63 kHz (Figure 4 E-H). Within the low load range of 0-100 Ω, the induced voltage remains relatively low, increasing linearly with the load. In the load range of 100-900 Ω, although the induced voltage continues to rise, the trend shifts, no longer adhering to linearity (Figure 4F). Beyond 900 Ω, the induced voltage nearly saturates (Figure 4E). This trend of induced voltage versus load was found to depend on the strength of the generated magnetic field in the transmitter coil. When a weaker magnetic field is generated in the transmitter circuit, the induced voltage saturates at a higher load, 6 kΩ; however, it was saturated at 1 kΩ at higher generated magnetic fields, 7 mT (Figure 4 G-H).

### Device Encapsulation

The receiver and electric circuits were assembled and encapsulated in a polymeric capsule. ICOPS performance was assessed through a series of benchtop tests. ICOPS is comprised of a coil wound around a nickel-zinc ferrite core, possessing an inductance of approximately 625.5 µH, a micro-capacitor, and the µLED. Two models were created: one resonated at 63 kHz, and the other at 100 kHz, utilizing capacitors of 10 and 4.7 nF, respectively.

The device was enclosed via a 3D-printed polymeric capsule. We used Veroultraclear resin which combines optical clarity with strength, stiffness, and impact resistance, making it suitable for ingestible electronic capsules.^28^

We printed the capsule using both vertical and horizontal printing orientations to maximize optical transparency. Both vertical and horizontal printing orientations were explored to maximize transparency and ensure robust mechanical properties. The capsule printed in a standing/vertical position on the printing bed exhibited greater transparency and durability compared to the horizontally printed one (Figure 5 A and B). The capsule walls were constructed with a thickness of 250 µm, and the total length of the capsule, including the lid, measured 18 mm. The inner diameter of the capsule was approximately 2.2 mm, while the outer diameter of the capsule body was around 2.7 mm, placing the capsule within the standard size range for rat-sized capsules. After incorporating the LED and electronics into the capsule, it was sealed with medical-grade epoxy (EPO-Tek-301). To assess the transparency of the polymeric matrix used, we printed a slab with a thickness of 250 µm in a vertical orientation, and measured its transmittance and absorbance across wavelengths (Figure 5C). The polymer slab exhibited a transmission peak at a wavelength of 500 nm, with a transmittance of approximately 80% for blue light of wavelength 460 nm, similar to the blue light emitted by the μLED connected to the encapsulated device.

**Figure 5.**
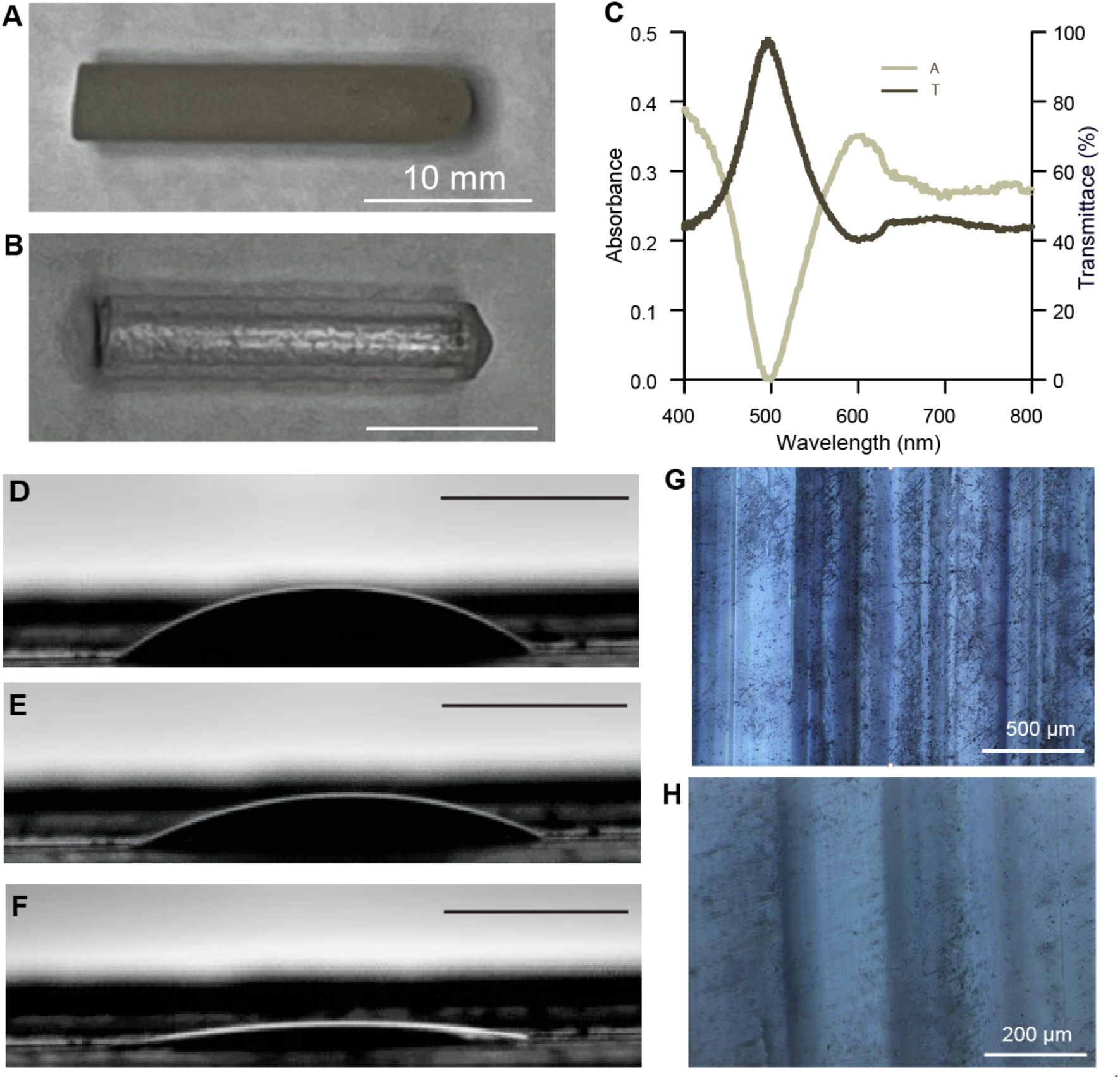
(A-B) Photographs of the polymeric capsules printed horizontally and vertically on the printing bed, respectively. (C) Transmission and absorption of the 3D-printed polymeric capsule. (D-F) Image of the 5 ml distilled water droplet on the 3D-printed slab recorded immediately upon pipetting, after 30 sec, and after a minute, scale bar: 10 mm. (G-H) Low and high magnification optical microscope images of the 3D-printed slab.

To better determine the interaction of the capsule surface with GI fluids, we evaluated the wettability by measuring the contact angle using the sessile drop method (Figure 5 D-F). Initially, the contact angle was measured at 40.6°, but it decreased over time as the droplet spread across the surface, reaching 30.1° within 30 seconds and 13.2° at 60 seconds (Figure 5 D-F). This reduction indicates a water-wicking effect on the ICOPS surface, which could help minimize the barrier between the ICOPS and the GI wall, allowing more light to penetrate and reach the targeted cells. This hydrophilic effect may be enhanced by microchannels formed during the layer-by-layer printing process, as observed on the surface under an optical microscope (Figure 5 G and H).^29^

### Benchtop Test of the Capsule

We next evaluated the performance of ICOPS using a benchtop setup. A transmitter with a diameter of 6.8 cm was tuned to 63 kHz, and the ICOPS and transmitter were arranged coaxially (Figure 6A). We moved the ICOPS inside the transmitter coil from one side of the wall to the other. As anticipated, the optical power emitted from the capsule exhibited a parabolic shape as the capsule traversed the coil (Figure 6B). The maximum optical power was detected at both walls of the coil, gradually diminishing to a minimum at the coil’s center. A similar parabolic shape was observed in the magnetic flux density measured along the diameter of the coil (Figure 6C). These magnetic field distribution measurements were supported by simulation results of magnetic field distribution inside the transmitter coil (Figure S3A).

**Figure 6.**
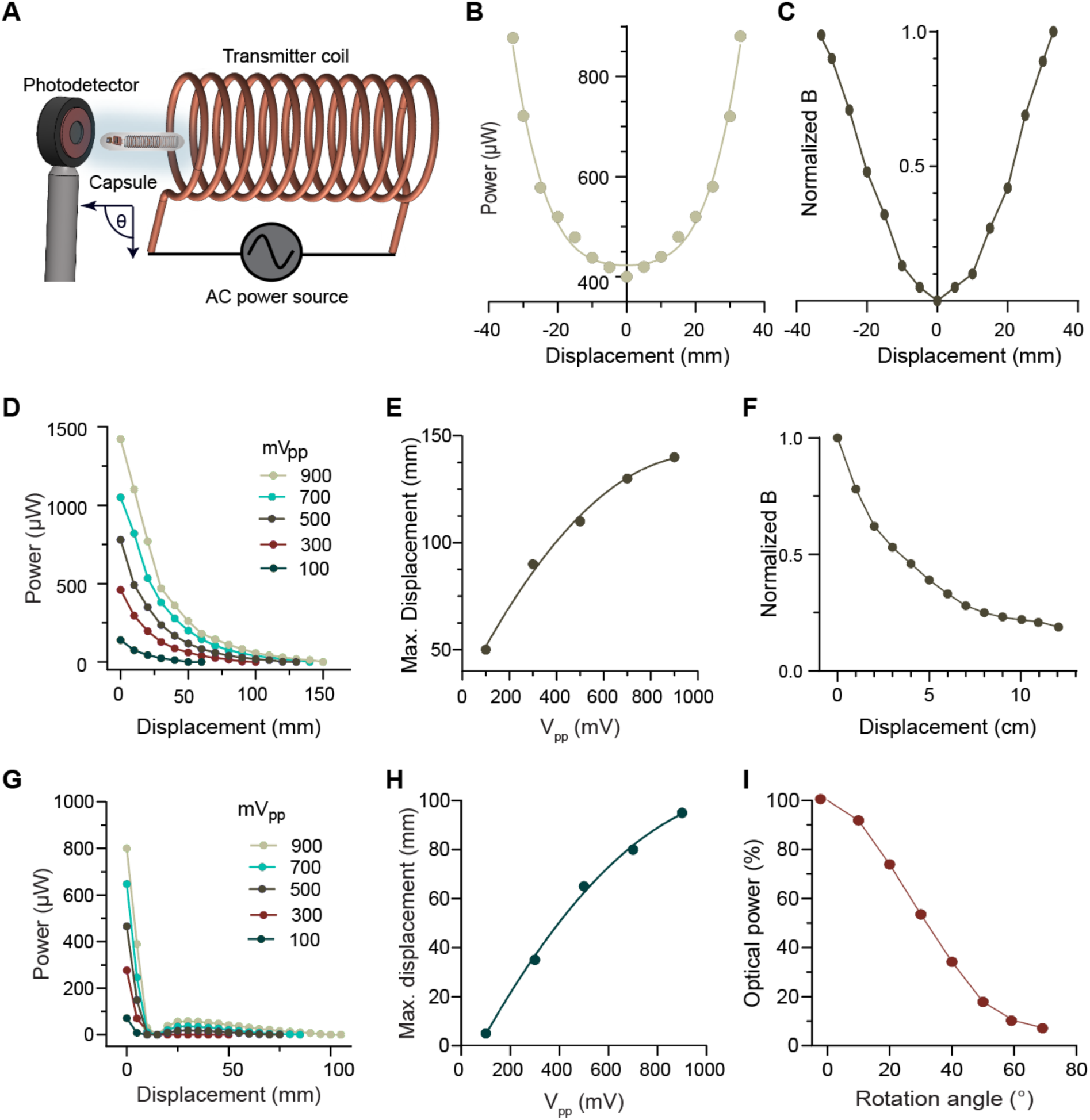
Benchtop tests of ICOPS performance. (A) Schematic illustrating the test setup used to record the ICOPS’s optical power under various conditions. (B) Output optical power of ICOPS across the transmitter coil’s cross-section, from edge to edge. (C) Normalized magnetic flux density along the diameter of the coil’s cross-section. (D-E) Effect of longitudinal displacement between ICOPS and transmitter on ICOPS optical power, demonstrating maximum operational displacement under different input powers in the transmitter circuit. (F) Trend of the magnetic flux density versus displacement from the transmitter. (G-H) Optical power of ICOPS versus lateral displacement from the transmitter under varying input powers in the transmitter circuit. (I) Optical power of ICOPS versus rotation angle relative to the magnetic field direction of the transmitter coil.

We next sought to assess the efficacy of wireless power transfer across ICOPS’ location and orientation. We quantified the optical power emitted by the ICOPS relative to its displacement from the transmitter, with both components aligned coaxially. We observed a nonlinear decline in optical power as the ICOPS distanced itself from the transmitter, a phenomenon in accord with the electromagnetic attenuation elucidated by the Beer-Lambert law (Figure 6D).^30^ Notably, a 50% reduction in output power occurred at a separation distance of 3 cm from the transmitter (approximately at a distance equals to the transmitter coil’s radius), diminishing from an initial 1400 μW to approximately 700 µW under the influence of a robust magnetic field engendered within the transmitter coil by an input voltage of 900 mV_p_. The maximum operational distance between ICOPS and the transmitter depends upon the potency of the magnetic field generated within the transmitter coil, with higher magnetic fields permitting extended distances. We found a maximum operational range of 12 cm in ambient air, corresponding to a magnetic flux density of approximately 7.2 mT, propelled by an applied voltage of 900 mV_p_ to the RF amplifier (Figure 6E). This operational range was reduced to less than 4 cm with a lower voltage input of 100 mV_p_ (Figure 6E). We measured the magnetic flux density across the same track where the capsule’s optical power was traced, and the *B* showed a similar trend to the measured optical power of the capsule (Figure 6F). As it decreased significantly to around 50% at 3 cm away from the transmitter which may explain the observed optical power trend of ICOPS. These external magnetic field measurements were supported by simulation results which showed the magnetic field nonlinearly dropped away from the transmitter coil showing a trend aligned well with the measured one (Figure S3B and S3C).

We next evaluated the optical power of ICOPS with lateral misalignment of 0-10 cm maintaining angular alignment throughout. Optical power attenuated to zero at lateral distances of 1-2 cm from the coil, yet increased with increasing distance, persisting up to 9 cm from the coil (Figure 6G). This long operation lateral distance correlated with the generation of a robust magnetic field (7.2 mT) within the coil, facilitated by the application of 900 mV_p_ to the RF amplifier. Lower applied voltages reduced the maximum operational lateral distance. The relationship between maximum operational lateral distance and applied voltage within the transmission circuit was found to be non-linear, possibly due to the non-linear RF amplifier and/or the non-linear I-V curve of the µLED (Figure 6H).

We examined the impact of angular misalignment of the capsule relative to the transmitter by systematically rotating ICOPS to varying angles, and measuring its optical power (Figure 6I). Even at an angular misalignment of 75°, the induced voltage within the receiver coil sufficed to energize ICOPS. The optical power of the capsule decreased to 10% at this degree of angular misalignment (Figure 6I).

We conducted an ingress test on the capsule to validate its integrity and ascertain its ability to endure the rigorous conditions within the GI tract. Three ICOPS were immersed in gastric fluid for a duration of 2 days and subsequently powered remotely upon removal from the fluid. All ICOPS were fully operational post-immersion and remained visually intact upon inspection (Figures 7 A and B, Figures S2 B and C). Before proceeding to ex vivo or in vivo tests, the capsule was examined using a µCT system to ensure it could be tracked after administration to a rat (Figure 7C). The device incorporated within the capsule was clearly visible and showed high contrast, making it easily detectable in the rodents due to its metal composition.

**Figure 7.**
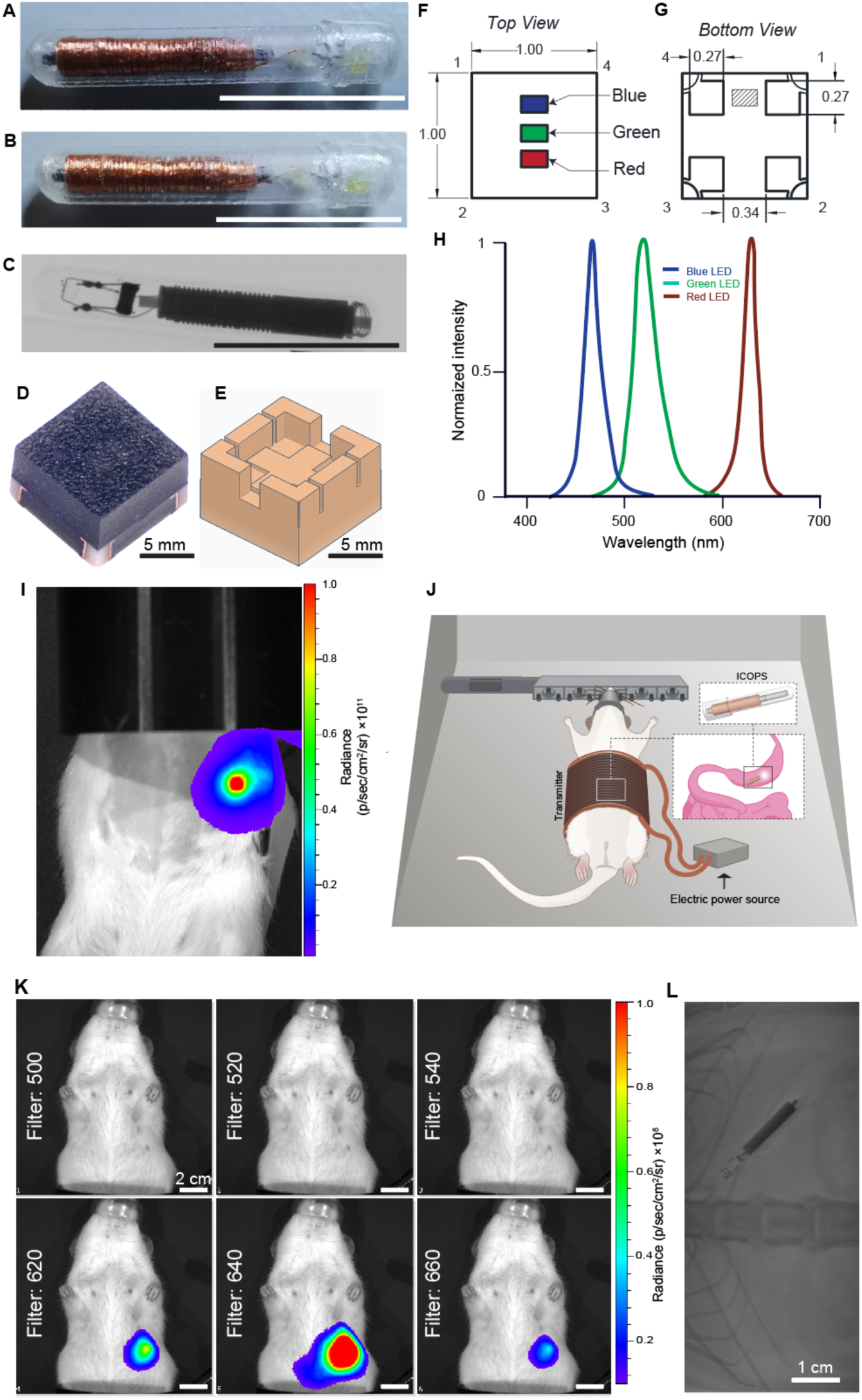
(A-B) ICOPS before (A) and after (B) the ingress test. (C) μCT image of ICOPS. (D) Photograph of the white LED. (E) Schematic of the 3D-printed mold designed for connecting the white LED. (F and G) Diagram illustrating the dimensions, top view, and bottom view of the white LED. (H) Light intensity of the white LED showing the dominant emitted wavelength from its embedded blue, green, and red LEDs. (I) IVIS image of ICOPS implanted in a sacrificed rat, equipped with a white light LED. (J) Schematic illustration of the ICOPS orally administered to a rat and wirelessly powered during imaging in the IVIS chamber. Scale bars 1cm unless otherwise noted. (K) IVIS image showing the rat administered with ICOPS and wirelessly powered, with light signals detected using various emission filters for blue, green, and red LEDs. (L) Two-dimensional μCT image of ICOPS after oral administration to the rat.

### Ex Vivo and In Vivo Tests of ICOPS

Initially, we tested ICOPS in an XFM-2 phantom mouse within the IVIS imaging system to assess the feasibility of monitoring its functionality within the rodent’s GI tract. We inserted ICOPS into the phantom and we remotely powered it to evaluate the potential detection of the emitted blue light signal. Proximity to the skin enabled the IVIS system to capture the blue light signal emitted from ICOPS (Figure S4A). However, as we inserted the ICOPS deeper into the phantom, signal detection became unattainable (Figure S4B). This attenuation phenomenon is well-documented, indicating a significant reduction in blue light transmission through tissue compared to light of longer wavelengths such as red light.^31^ To address this issue, we decided to include a red LED in the ICOPS to indicate its functionality.

A milli-scale red LED emitting at around 630 nm was powered and positioned underneath the mouse phantom in the IVIS chamber. The IVIS system successfully captured the light signal, demonstrating that the red light of that wavelength easily penetrated the entire mouse phantom tissue, producing a strong signal at the phantom’s skin surface (Figure S4C). The red light may serve dual functions: monitoring the functionality of the capsule and activating/silencing red channelrhodopsin.^32^ We also incorporated a green μLED, allowing ICOPS to activate/silence blue, green, and red channelrhodopsin.

We used a white LED composed of blue, green, and red µLEDs, with the blue µLED emitting at a wavelength of 470 nm, the green µLED emitting at 526 nm, and the red µLED emitting at 622 nm (Figure 7 D-H). The white LED-equipped capsule was then evaluated in a sacrificed rat following implantation into the stomach. Upon remote powering, the IVIS imaging system successfully detected a robust signal from ICOPS (Figure 7I). With the feasibility of ex vivo functionality monitoring established, we orally administered the capsule to an anesthetized rat using a modified gavage tube (Figure 7J). Next, we placed the rat inside the IVIS Spectrum for optical imaging while the rat was under anesthesia. Subsequent wireless powering of ICOPS, a series of images were acquired using different emission filters to distinguish the wavelength of the emitted light reaching the camera (Figure 7K). The signal emitted in the far-red window which corresponds to the red LED was detected, confirming the superior penetration capability of the red light compared to blue and green. Additionally, μCT was used to confirm the location of the administered ICOPS. The high distinguishability of the capsule’s metal coil from the surrounding rat tissue ensured easy detection (Figure 7L, Figure S5 A-E).

### Specific Absorption Rate

Modern RF exposure standards are primarily expressed in terms of specific absorption rate (SAR), which is defined as the amount of power dissipated per unit mass. The International Commission on Non-Ionizing Radiation Protection (ICNIRP, 1998) sets a SAR limit of 0.4 W/kg, averaged over the whole body, for continuous and occupational exposure. ^21,33^ This applies to informed, healthy adults.

The transmitter coil was modeled using Ansys Electronic Desktop 2023R1, with a rat trunk phantom approximated as a cylinder with a radius of 32 mm and a height of 80 mm. The relative permittivity of the rat trunk was assumed to be 100, and the conductivity was taken as 0.5 S/m, which is the average value for human bones, muscles, and tissues as reported in the literature.^34^ The 5/20/38 AWG wire used in the experiment, with an overall diameter of 1.32 mm, was approximated as a conductor with a diameter of 1.01 mm and a total insulation layer diameter of 0.31 mm. Table S1 summarizes the design considerations.

In the SAR simulation, the maximum current that could be driven in the coil by our 50 W RF amplifier was considered, which is 6 A (rms). The maximum SAR value at the operating frequency of 45 and 63 kHz was found to be 0.14 and 0.26 W/kg, respectively, which are below the occupational exposure safe limit, indicating that the power transfer system is unlikely to have a significant impact on the core body temperature (Figure S6). However, at higher operating frequencies of 81, 100, and 140 kHz, the SAR values exceeded the safe limit for continuous exposure.

## Discussion

Efforts to adapt standard wireless implantable optical stimulators, originally designed for the brain and spinal cord, for application in the GI tract, have underscored the necessity for technologies tailored specifically to this anatomical region. Previous studies have highlighted challenges; for instance, optogenetic stimulation of ex vivo whole colon using these devices resulted in contractile responses of the musculature in only 13 out of 20 trials (n = 5), even when applied over a partial length of the colon known as colonic motor complexes (CMCs). Moreover, when surgically implanted in vivo, these devices induced colonic obstructions in 60% of the tested mice.^17^ These implantable devices rely on passive electronic circuits constructed using semiconductor microfabrication techniques, necessitating clean room facilities for manufacturing. This requirement prolongs and complicates the production process, demanding extensive training. Furthermore, any modifications to the device design require significant efforts in adjusting and optimizing numerous process steps, resulting in prolonged production times. Additionally, they face challenges in fabricating intricate three-dimensional (3D) shapes and features to enhance their functionality. Traditional silica fibers have been found to puncture the mucosal membrane during intraluminal implantation in the mouse small intestine. Additionally, surgical implantation often requires bending the fiber at acute angles, which significantly reduces the delivered irradiance. As a result, traditional fibers are not suitable for in vivo use in the gut.^35^

Addressing inherent limitations associated with implantable optical stimulators, such as susceptibility to infection, inflammation, or rejection of the implanted foreign body, is crucial. These challenges prompted the development of our novel system: a passive electronic circuit housed within an ingestible electronic capsule powered wirelessly. ICOPS offers several advantages over conventional devices: it is noninvasive, enhancing animal comfort and avoiding tissue damage; it does not require semiconductor microfabrication techniques; it operates as a multi-point stimulation system capable of withstanding peristaltic motion and the harsh GI tract environment without needing to connect with the GI tract walls; and it can activate or silence channelrhodopsin variants in the visible light range, including blue, green, and red.

This proof-of-concept study presents the ingestible electronic capsule designed for optical stimulation of the GI tract. The capsule-integrated LEDs emit sufficient blue light to activate or silence channelrhodopsin2, and the resonance capacitance technique efficiently resonates the transmitter and capsule at the operating frequencies. Notably, the capsule can operate with less than 1% of the amplifier output power (500 mW). ICOPS functions effectively up to 12 cm away from the transmitter. Both ex vivo and in vivo testing confirm the capsule’s functionality within the GI tract, with the added benefit of a red LED facilitating monitoring using the IVIS system. The incorporation of green LED alongside blue and red LEDs enables comprehensive activation or silencing of channelrhodopsin across the visible light spectrum. Furthermore, the capsule’s metal components are easily distinguishable from surrounding rat tissue, allowing for straightforward detection after ingestion using μCT.

Future iterations of ICOPS should address technological concerns to enable optical stimulation in freely moving animals. Redesigning the transmitter coil into a wearable jumper for rats would facilitate optical stimulation while animals remain conscious and mobile. Continued advancements in wireless technology could lead to miniaturization of the current cabled wireless transmission system, enhancing portability. Moreover, integrating additional LEDs into the capsule could increase irradiance, improving the feasibility of optical stimulation even when food acts as a barrier between the capsule and GI tract walls. Given the dimensions of the receiver coil, capacitor, and LED holders, there is ample space to accommodate these enhancements while maintaining the capsule within standard size ranges.

## Conclusion

The ICOPS is a battery-free, untethered, orally administered capsule for optical illumination within the GIT. With the growing focus on understanding gut-brain axis communication in various health and disease states, ICOPS could be a potential tool for non-invasive optogenetic manipulation of enteric neural circuits. An improved understanding of these circuits could pave the way for novel therapies leveraging enteric neurons and the gut-brain axis.

## Materials and Methods

### Optoelectronic Characterization of the µLED

To examine the I-V characteristic curve of the µLED (Cree^®^TR2432^TM^), it was linked to a DC power supply, and the current was documented against the applied voltage.

Experimental procedures to measure the optical power of the utilized μLED involved connecting it to the DC power supply. A photodiode (Thorlabs, PM16-120) was employed to capture the optical power corresponding to the applied voltage. The identical configuration was employed to ascertain the dominant wavelength of the spectrum emitted from the µLED, except a spectrometer (Ocean Insight, USB 2000+) was utilized instead of the photodiode to gauge light intensity.

### Fabrication of the Capsule (ICOPS)

A stereolithography-based 3D printing system (BMF microArch^®^ S240), offering a resolution of 10 µm, was employed to fabricate molds for both the µLED and the micro-capacitor integrated into the capsule (**Figure S2A**). These molds were designed using Tinkercad, an accessible online software. The designs were saved in .stl format, then sliced using the BMF printer software, and subsequently printed layer-by-layer. The printing duration varied depending on the number of sliced layers and the UV exposure time, with an average printing time of approximately 2 hours. The µLED and capacitor were affixed within the molds using low-temperature soldering epoxy (Chip Quik, TS391LT10) and subsequently connected in parallel with the receiver. The receiver coil was wound around a nickel-zinc ferrite core with a diameter of 0.675 mm. This coil comprises 5 layers, totaling 460 turns, and was constructed from copper wire with a diameter of 125 µm, including the insulation layer, resulting in an external diameter of 2.1 mm. The inductance of the coil was measured to be 625.5 µH within the frequency range of 45-140 kHz using an LCR meter (Hioki, IM3536).

The assembled device was encapsulated within a polymeric capsule fabricated using the Stratasys J750 3D printing system. The capsule was designed on the Tinkercad Webpage and printed with VeroUltraClear resin. Printing was conducted in two orientations: vertically (standing up) and horizontally (lying down). The capsule walls had a thickness of approximately 250 µm. The evaluation of the optical transparency of the encapsulating polymer was conducted through measurements of its transmittance and absorbance. Utilizing a square slab measuring 10 mm on each side, the polymer was printed and subjected to investigation employing a spectrometer (Ocean Insight, USB 2000+). The polymer’s wettability was scrutinized by employing the sessile drop method (Dataphysics Instruments, OCA 15EC). This involved the precise deposition of a 5 ml droplet onto the printed polymer slabs, with subsequent capture of temporal images of the droplets. The surface morphology of the printed slabs was examined utilizing an optical microscope (Nikon LV-Dia) in bright field mode employing both 5x and 10x microscope lenses.

### Tuning the Transmitter and Receiver Circuits

The transmitter coil was fabricated using an 18-gauge Litz wire wound around a hollow plastic cylinder with 5 mm thick walls, manufactured via 3D printing using an UltiMaker printer (UltiMaker 2+ Connect). Its dimensions include a diameter of 68 mm and a length of 70 mm, with 50 turns forming a single layer. The coil’s inductance was quantified through measurement using an LCR meter. Driving the transmitter coil involved a function generator (Siglent, SDG 2042X) interfaced with an amplifier (Accel Instruments, TS250) with a maximum output power of 50 W and a maximum current of 6 A (rms). Initially, the resonance of the transmitter coil circuit was achieved by incorporating a capacitor in series and sweeping the frequency range to ascertain the resonance frequency, concurrent with the measurement of magnetic flux density using a Gaussmeter (Hirst Magnetic instruments, GM07). Subsequently, the alignment of the receiver circuit to the same operational frequency was ensured by introducing a micro-scale shunt capacitor. At this designated operational frequency, capacitance variation ensued, with measurement of the optical power output from the µLED conducted via the power meter. Selection of the capacitor optimizing optical power marked resonance of the receiver circuit. The capsule’s performance underwent evaluation after successful resonance establishment in both transmitter and receiver circuits at the specified operational frequency range. Specifications of both the transmitter and receiver coils are summarized in Table 1 below.

**Table 1.** Specification of the receiver and transmitter coils.

| Specs | Transmitter coil | Receiver coil |
| --- | --- | --- |
| Diameter(cm) | 7 | 0.21 |
| Length (cm) | 6.8 | 1 |
| Thickness (mm) | 1 | 0.7 |
| Inductance (μH) | 122.55 | 626.5 |
| Parasitic capacitance (pF) | 3.8 | - |
| Self-resonance frequency (MHz) | 3.57 | 14.67 |
| Mutual inductance (μH) | 0.499 | 0.499 |
| Coupling coefficient | 0.0018 | 0.0018 |

### Benchtop Test of the Capsule

The output power of the capsule was assessed as it moved along the diameter of the transmitter’s coil cross-section. Measurements were taken at 5 mm intervals while the capsule, along with the photodetector, remained fixed to a moving stage. This stage traversed from one wall of the coil to the opposite wall, passing through the coil’s central point. Concurrently, magnetic flux density was measured at 5 mm intervals along the same trajectory using the gaussmeter. Furthermore, the output power of the capsule was evaluated concerning its displacement from the edge of the transmitter coil along its long axis, with both the receiver and transmitter’s long axes being parallel. Additionally, the capsule’s output power versus displacement was recorded while the capsule moved perpendicular to the long axis of the transmitter coil. Finally, the output power of the capsule was examined concerning its orientation angle relative to the long axis of the transmitter coil/generated magnetic field.

### Ex Vivo and In Vivo Testing of ICOPS

All animal procedures were approved by the Animal Care and Use Committee of New York University Abu Dhabi, and protocols were conducted in compliance with the guidelines outlined in the National Institute of Health Guide for Care and Use of Laboratory Animals (IACUC protocol number: 21-0007A4). Female rats aged 12 months were utilized (Sprague Dawley rat strain). The rats were provided with food and water ad libitum and maintained under a 12-hour light-dark cycle. The functionality of ICOPS was assessed through various methods. First, ICOPS was inserted into a hole at the center of a mouse phantom, and the emitted blue light signal was detected using the IVIS imaging system (Perkin Elmer, USA). Second, ICOPS was tested in a euthanized rat, implanted in the stomach, and its functionality was evaluated using IVIS. Prior to ICOPS testing, the rat was euthanized using a CO_2_ chamber Third, the capsule’s functionality was observed in a live rat under anesthesia. The rat was anesthetized with isoflurane, induced at 5%, and then maintained at 2% in a 1 L/min oxygen flow. The depth of anesthesia was monitored by assessing the absence of response to a hind limb or tail pinch. The rat was positioned within the IVIS, which has an adjustable temperature chamber set to 37°C. After assessing ICOPS functionality via IVIS, the administered ICOPS was tracked using µCT imaging (Skyscan µCT, Bruker, USA).

