## Supplemental Information for "Wirelessly-Powered Ingestible Electronic Capsule for Non-invasive Gastrointestinal Optogenetics"

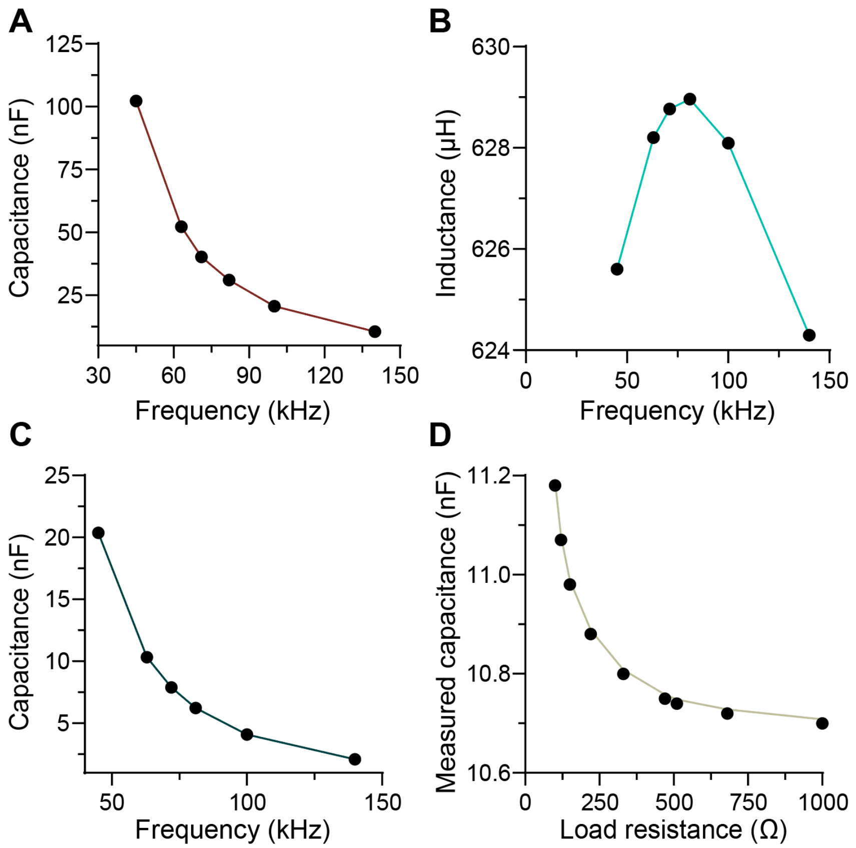


**Figure S1.** (a) Measured parasitic capacitance of the transmitter coil across the operating frequency range (45-140 kHz). (B) The inductance of the receiver coil across the operating frequency range. (C) Measured parasitic capacitance of the receiver coil across the operating frequency. (D) Measured capacitance of the receiver circuit (receiver coil connected in parallel with various load resistances) at different load resistances measured at 63 kHz.

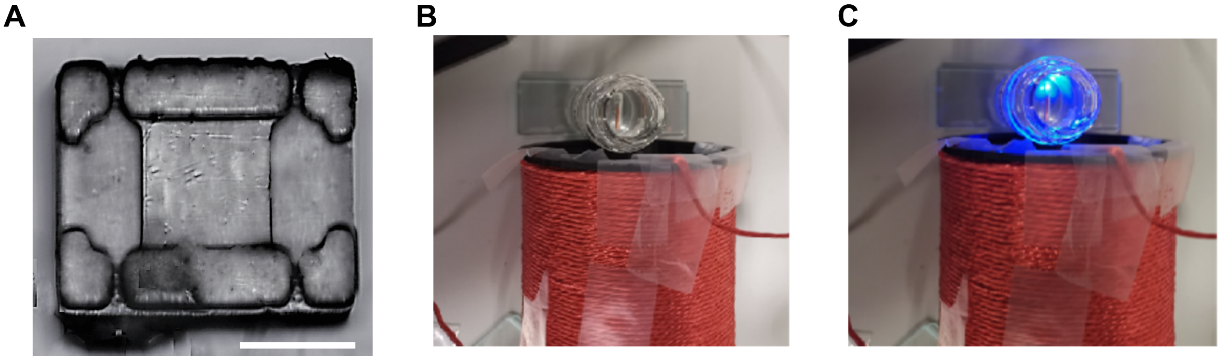


**Figure S2.** (A) Optical microscope image of the mold used to connect the used micro-capacitor, scale bar: 500μm. (B and C) Photographs of the capsule immersed in gastric fluid in ON and OFF conditions, respectively.

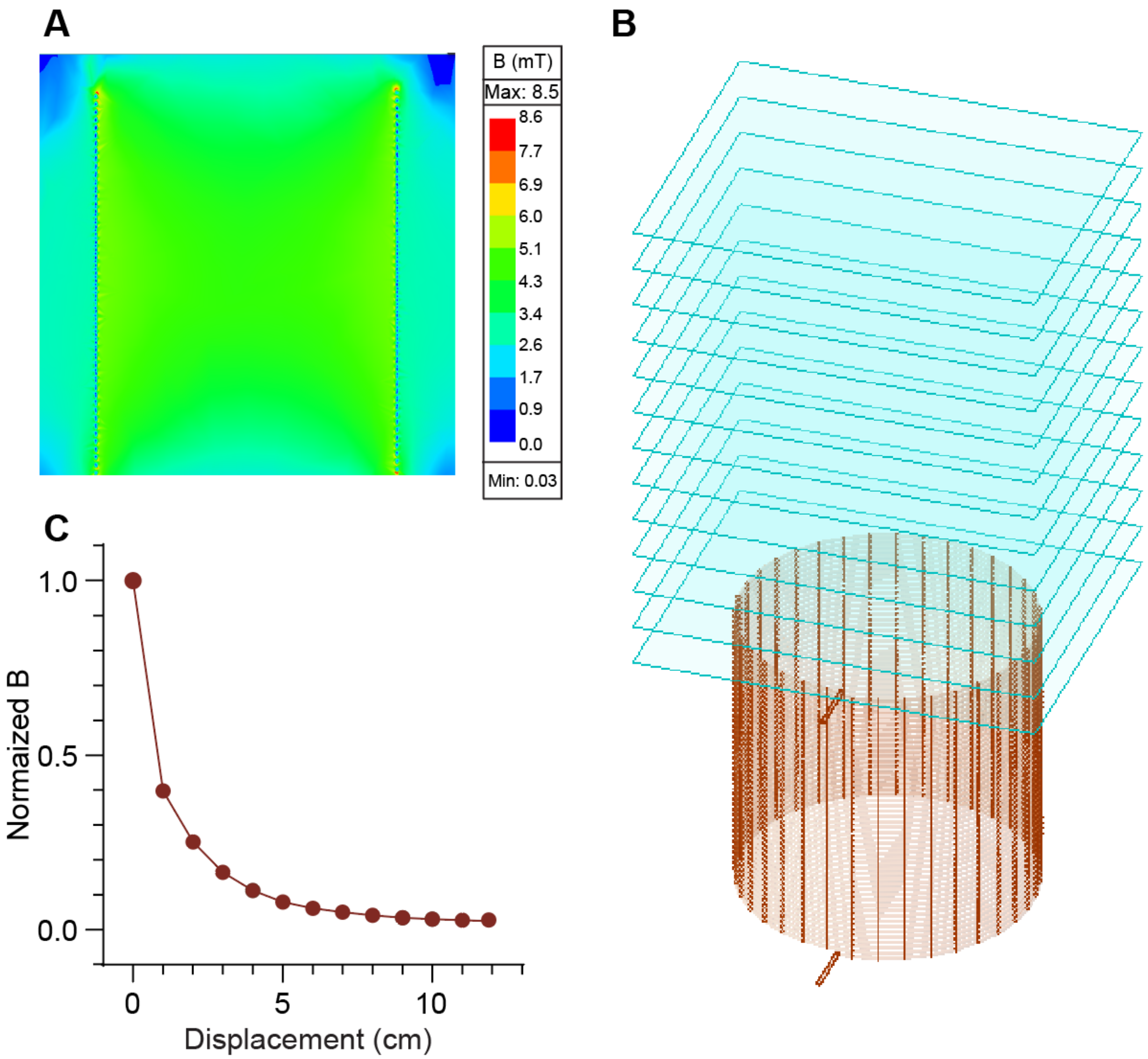


**Figure S3.** (A) Magnetic field distribution inside the transmitter coil. (B) Schematic depicting the transmitter coil and the simulated planes considered for calculating the magnetic field density. (C) Normalized magnetic flux density plotted against displacement from the coil.

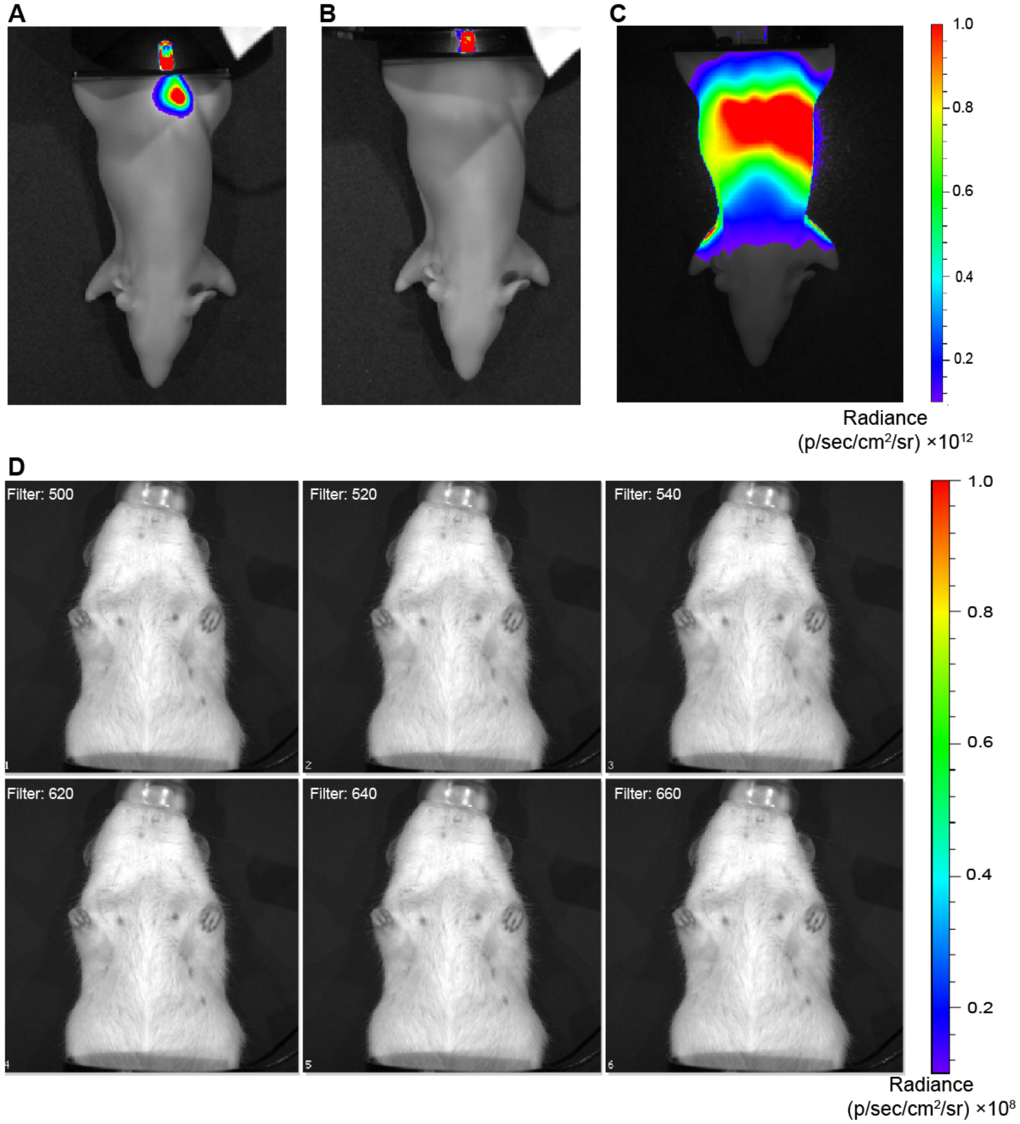


**Figure S4.** (A-B) IVIS images of a mouse phantom with the capsule inserted in the middle and deeper holes of the body, respectively. (C) IVIS image of a mouse phantom with a red LED positioned underneath. (D) IVIS images of an anesthetized rat orally administered with the capsule equipped with blue, green, and red LEDs in the Off condition, different filters are applied.

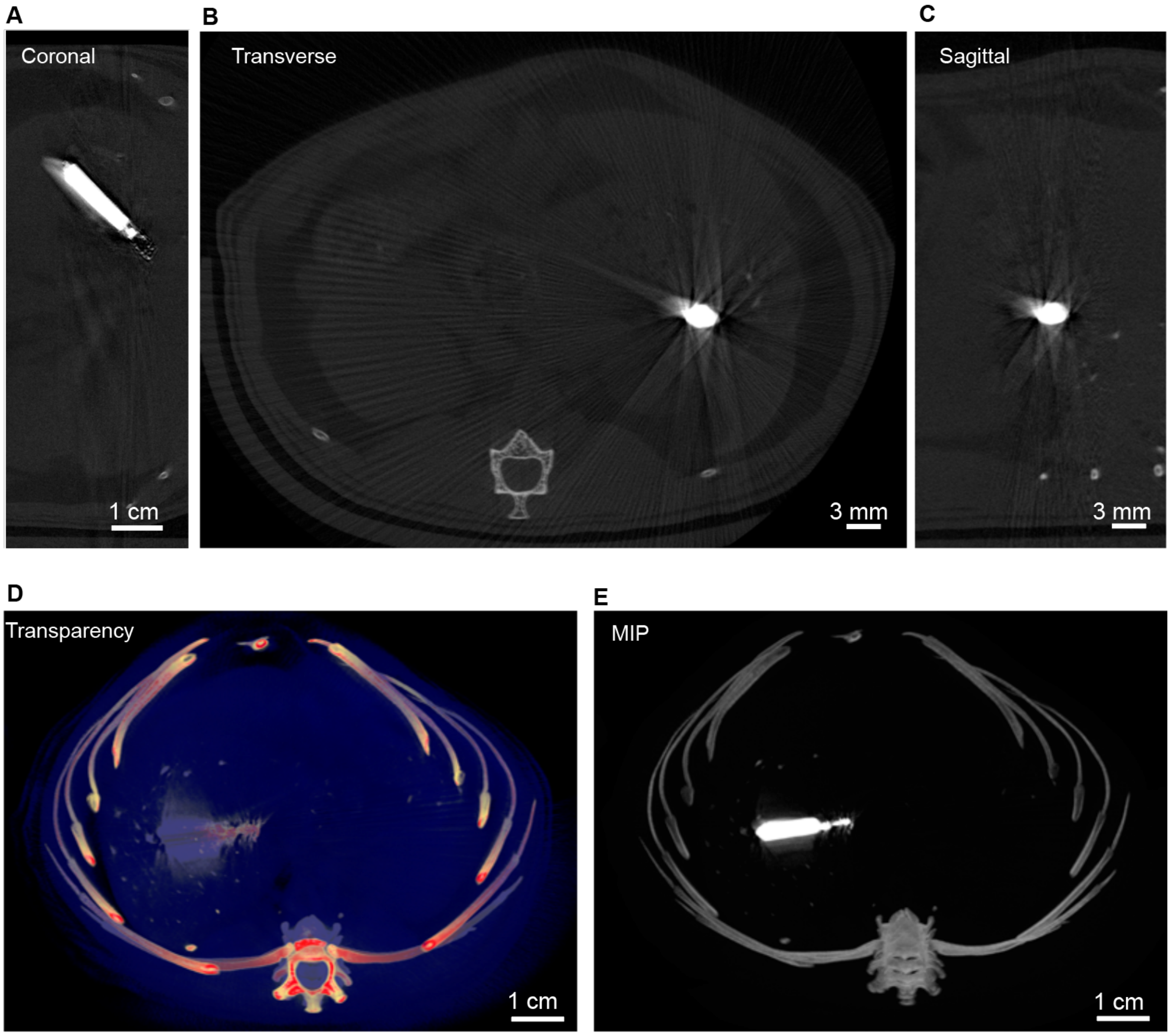


**Figure S5.** Three-dimensional μCT images of ICOPS after oral administration to a rat.

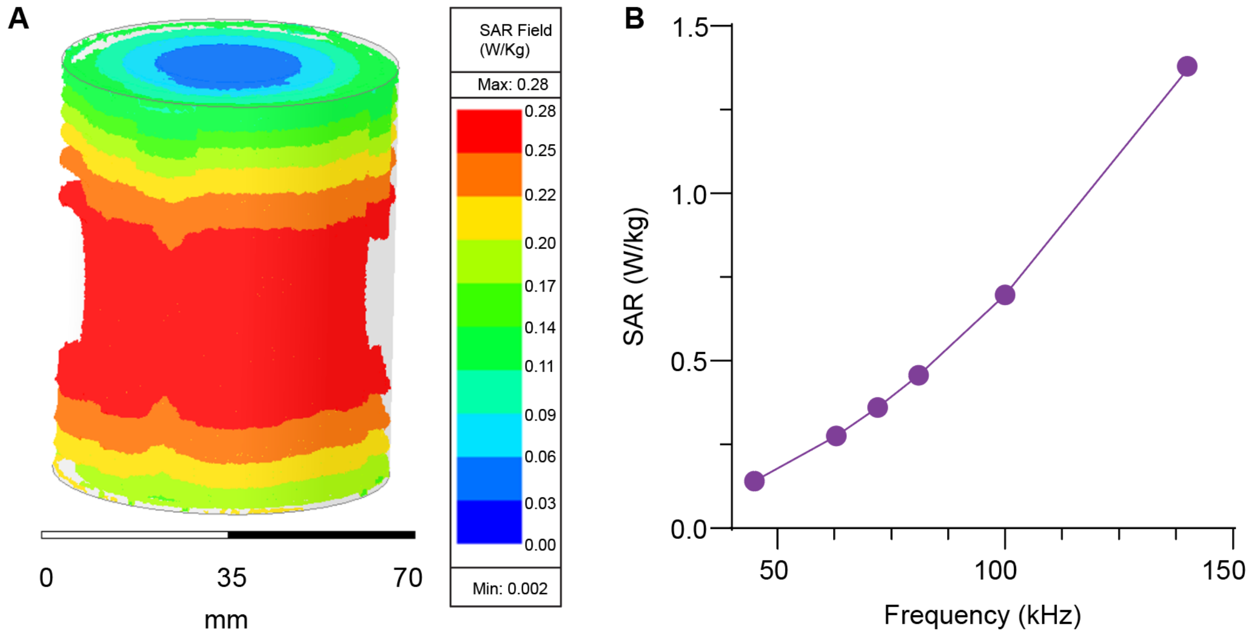


**Figure S6.** Specific absorption rate (SAR) in body tissues. (A) SAR of body tissues exposed to the maximum magnetic field generated by the transmitter at a driven current of 6 A (rms) operated at a frequency of 63 kHz. (B) SAR of body tissues exposed to maximum magnetic fields generated by the transmitter at various frequencies (45-140 kHz) with a maximum current of 6 A (rms) in the coil.

**Table S1.** Transmitter coil design consideration for simulating the SAR

| Design Variable | Value |
|---|---|
| Length of the coil | 73 mm |
| Diameter of the coil | 73. 4 mm |
| Number of turns | 50 |
| Equivalent diameter of the conductive wire | 1.007 mm |
| Spacing between the wires | 0.459 mm |
| Wire material conductivity | 52426000 S/m |
| Rat phantom cylinder radius | 32 mm |
| Rat phantom cylinder height | 80 mm |
| Rat phantom material permittivity | 100 |
| Rat phantom material conductivity | 0.5 S/m |
